# Evolution of the quorum sensing regulon in cooperating populations of *Pseudomonas aeruginosa*

**DOI:** 10.1101/2021.07.01.450773

**Authors:** Nicole E. Smalley, Amy L. Schaefer, Kyle L. Asfahl, Crystal Perez, E. Peter Greenberg, Ajai A. Dandekar

**Affiliations:** Department of Medicine, University of Washington, Seattle, WA 98195; Department of Microbiology, University of Washington, Seattle, WA 98195

**Author notes:** Corresponding author: Ajai A. Dandekar, K-359A HSB, UW Box 356522, 1705 NE Pacific St. Seattle, WA 98195.

**Keywords:** Acylhomoserine lactone, adaptive evolution, metatranscriptomics, quorum quenching

## Abstract

The bacterium *Pseudomonas aeruginosa* is an opportunistic pathogen and it thrives in many different saprophytic habitats. In this bacterium acyl-homoserine lactone quorum sensing (QS) can activate expression of over 100 genes, many of which code for extracellular products. *P. aeruginosa* has become a model for studies of cell-cell communication and coordination of cooperative activities. We hypothesized that long-term growth of bacteria under conditions where only limited QS-controlled functions were required would result in a reduction in the size of the QS-controlled regulon. To test this hypothesis, we grew *P. aeruginosa* for about 1000 generations in a condition in which expression of QS-activated genes is required for growth. We compared the QS regulons of populations after about 35 generations to those after about 1000 generations in two independent lineages by using quorum quenching and RNA-seq technology. In one evolved lineage the number of QS-activated genes identified was reduced by about 70% and in the other by about 45%. Our results lend important insights about the variations in the number of QS-activated genes reported for different bacterial strains and, more broadly, about the environmental histories of *P. aeruginosa*.

## Introduction

The cell-density-dependent activation of certain genes in the opportunistic pathogenic bacterium *Pseudomonas aeruginosa* relies in part on acyl-homoserine lactone (AHL) quorum sensing (QS). There are two *P. aeruginosa* AHL circuits, the Las and Rhl circuits. The genes for these circuits are conserved among hundreds of *P. aeruginosa* isolates from different environments. The Las circuit relies on the product of the *lasI* gene, an AHL synthase, for production of the diffusible QS signal *N*-3-oxo-dodecanoyl-homoserine lactone (3OC12-HSL), and on the product of the *lasR* gene, a transcription factor, which activates dozens of genes in response to 3OC12-HSL. The product of the *rhlI* gene is a butanoyl-homoserine lactone (C4-HSL) synthase, and the product of the *rhlR* gene is a C4-HSL-responsive transcription factor, which activates a set of genes that overlaps with those activated by LasR (Schuster et al. 2013). In PAO1 and many other isolates the Rhl circuit requires activation by the Las circuit. One isolate, *P. aeruginosa* PAO1, is particularly easy to manipulate genetically and the genome of this strain was the first *P. aeruginosa* genome sequenced (Stover et al. 2000). Strain PAO1 has become a model for studies of QS and for sociomicrobiology investigations (Schuster et al. 2013).

The genes activated by QS in strain PAO1 have been identified by comparing QS mutants to their parent (Whiteley et al. 1999, Schuster et al. 2003, Wagner et al. 2003, Chugani et al. 2012). Transcriptome comparisons have shown that well over 100 genes are activated by QS in this strain (Schuster et al. 2003, Wagner et al. 2003). Less is known about QS-dependent genes in other isolates, but transcriptomics of QS mutants of several isolates from different environments revealed the number of genes in the quorum regulon can vary from a few dozen to over 300, depending on the isolate. There was a core set of 42 genes activated by QS in most of the studied isolates (Chugani et al. 2012).

Many, but not all, QS-activated genes code for extracellular proteins or production of extracellular products. This is consistent with the idea that QS serves as a cell-cell communication system, which coordinates cooperative activities such that cells produce extracellular factors only when they have reached sufficient population densities (Fuqua et al. 1994, Schuster et al. 2013). This hypothesis has been tested in several ways. For example, a prediction of the hypothesis is that, because there is a fitness cost to cooperation, QS mutants should have a fitness advantage over their cooperating parent. This is in fact the case. Growth on proteins like casein or bovine serum albumin relies on QS activation of genes coding for extracellular proteases, and LasR mutants have a fitness advantage over their parent during co-culture on these proteins as carbon and energy sources (Diggle et al. 2007, Sandoz et al. 2007). Furthermore, when *P. aeruginosa* is cultured continuously with casein as the sole source of carbon and energy, LasR mutants emerge and reach a substantial fraction of the population within 10-20 days of transfer (Sandoz et al. 2007, Dandekar et al. 2012).

Not all QS-activated genes code for extracellular products and there is some understanding of the significance of this fact. One such gene, *nuh*, encodes a nucleoside hydrolase required for growth on adenosine (Heurlier et al. 2005). Strain PAO1 grows slowly on adenosine but fast-growing QS-dependent variants with amplification of a genomic region including *nuh* emerge upon sustained growth on adenosine (Toussaint et al. 2017). The addition of adenosine as a second carbon and energy source in addition to casein constrains the emergence of LasR mutants of strain PAO1 (Dandekar et al. 2012).

We are interested in understanding how QS circuits might evolve in *P. aeruginosa* adapted to life in different environments. Might such understanding help us understand the observed variation in QS regulons of different *P. aeruginosa* isolates and reveal something about the life history of any given isolate? The hypothesis underlying our investigation is that continuous growth of *P. aeruginosa* in an environment requiring activation of a few QS-activated genes will result in a reduction in the number of genes activated by QS – i.e., there will be a fitness advantage for individuals that do not activate genes of no benefit in the environment. Because LasR mutants arise within days of *P. aeruginosa* growth on casein, we provided *P. aeruginosa* with both casein and adenosine in our experiments. Growth on casein involves production of excreted proteases, which are considered as public goods. Growth on adenosine requires a cell-associated nucleosidase, which is considered as a private good. As discussed above, when adenosine and casein are both provided to the bacteria, LasR mutants are constrained for at least 30 days of daily transfer (Dandekar et al. 2012). Our experiments involved daily culture transfer for 160 days during which time we expect that a myriad of variants with different SNPs will arise. Thus, our QS regulon analysis was performed on populations rather than individual isolates. Our work therefore required development of a transcriptome comparison that did not involve a comparison of QS mutants to parents: a metatranscriptome analysis of QS gene activation.

## Results

### Long-term growth experiments

*P. aeruginosa* cells grown in buffered Lysogeny Broth (LB) were used to inoculate five tubes of a minimal broth containing only casein and adenosine (CAB) as carbon and energy sources. After inoculation, cells were transferred to fresh CAB daily for the next 160 days (Figure 1A). A 30-day experiment showed that QS mutants are constrained in CAB (Dandekar et al. 2012). A previous publication has shown that after the first several days of growth on casein, a PsdR mutant sweeps through the population. Mutations in *psdR* enhance growth on casein (Asfahl et al. 2015). The *psdR* gene codes for a repressor of a peptide transporter, and enhanced growth can be explained, at least in part, by improved uptake of peptide products of LasR-induced extracellular proteases. Growth on adenosine (or casein and adenosine) results in emergence of cells which show improved growth on adenosine resulting from a genome amplification of a region containing the LasR-dependent *nuh* gene (Toussaint et al. 2017). The gene-amplification variants retain LasR-dependent growth on adenosine.

**Figure 1.**
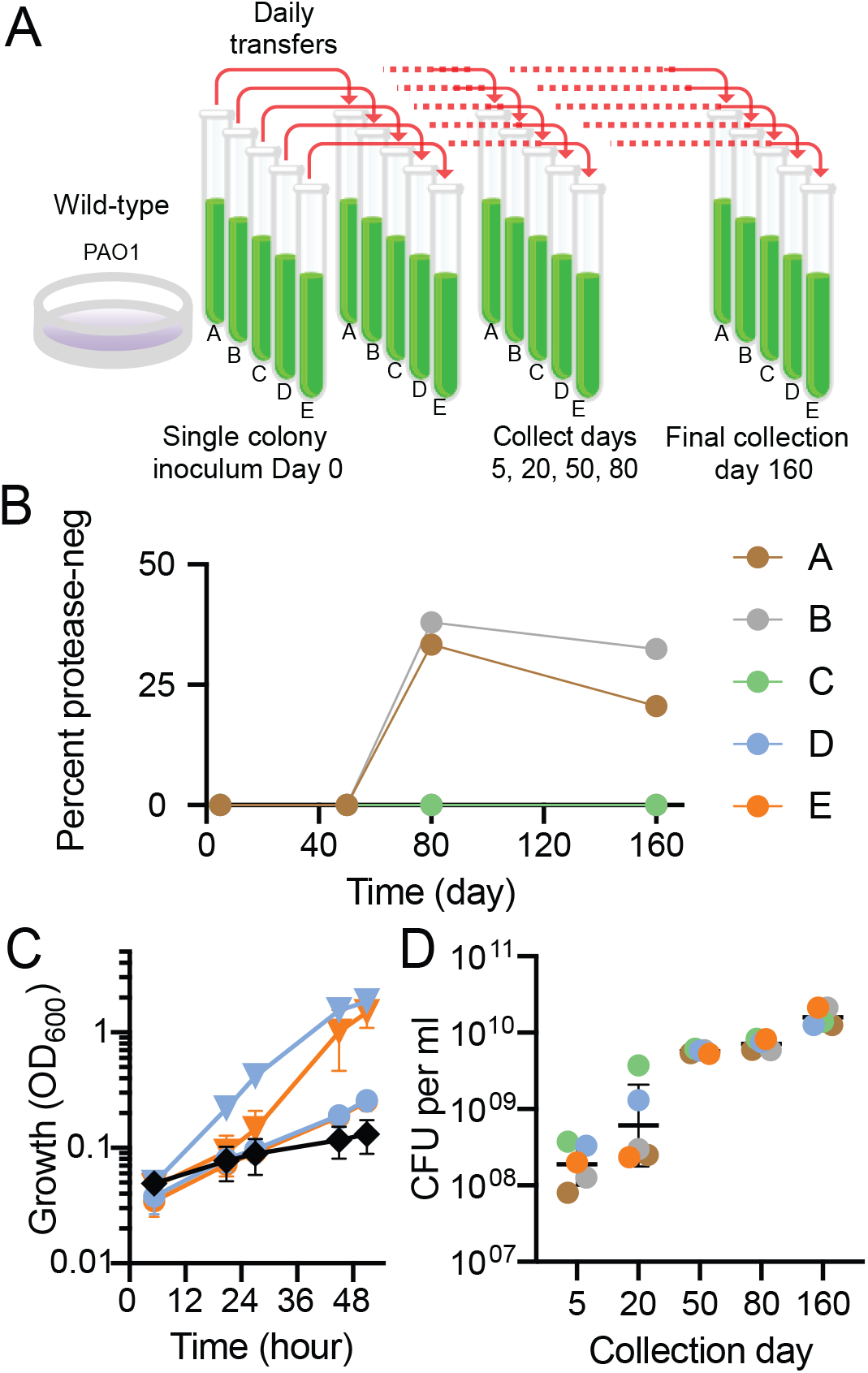
Long-term evolution of *P. aeruginosa* PAO1 serially passaged in a medium (CAB) that requires quorum sensing for growth. (A) Experimental design. (B) Abundance of protease-negative (neg) cheaters in each lineage (A thru E) at the indicated days of serial passage in CAB. (C) Growth in adenosine-only (1% weight/vol) broth of lineages D (blue) or E (orange) collected after 5 days (circles) or 50 days (triangles) of serial passage in CAB. Growth of the parent strain PAO1 (black diamonds) is included for comparison. Data are the means of two biological replicates, error bars are the range. (D) Growth yields of indicated populations grown in CAB for 18 h as determined by colony forming units (CFU) per ml. Black lines are the geometric mean of three or four biological replicates for each population; error bars are the geometric standard deviation.

Protease-negative isolates from growth in casein are almost always LasR mutants (Sandoz et al. 2007). We screened populations from 5-day, 50-day, 80-day and 160-day cultures of all five lineages for protease-negative individuals (Sandoz et al. 2007). We did not observe any protease-negatives among the 100 individuals from each of the 5-day or 50-day cultures, but protease-negatives were identified in two of the 80- and 160-day populations (Figure 1B). One likely explanation for breakthrough of protease-negative mutants after 160 days is that there is a mutation uncoupling adenosine metabolism from the QS-regulon such that LasR mutants do not incur a cost in CAB; they have a fitness advantage identical to that observed during growth of *P. aeruginosa* on casein alone.

We selected two of the three lineages (lineages D and E), in which protease-negative mutants were not detected, for further study. Early in the experiment at day 5, *psdR* mutations were not detected in either lineage D or E. At day 50 specific *psdR* mutations were fixed in each population. There was a frameshift mutation in lineage D (insertion of a C at position 431), and a T515C SNP in lineage E. This finding was consistent with the idea that the populations were adapting to growth on casein. We asked if an analogous adaptation was occurring for adenosine. When transferred to a minimal medium containing only adenosine as a carbon and energy source, day-5 populations of both lineages grew slowly, but day-50 populations grew more rapidly (Figure 1C), as would be expected of a population with a *nuh* region amplification. A next question is how does evolution in CAB affect population growth? Because partial degradation of casein results in a milky haze in CAB, growth cannot be monitored by following optical density. Therefore, we assessed growth yields by using plate counting to determine cell yields after 18 h in CAB for populations isolated after 5, 20, 50, 80 and 160 days, and found that there was a large increase in yield at day 50 (Figure 1D). This corresponded to a time after *psdR* mutations had swept through the two populations, and the populations had acquired the ability to grow rapidly on adenosine (Figure 1C). Because cultures were diluted into fresh CAB daily as a 1% inoculum there were six to seven doublings per day, corresponding to about 1,000 generations over the 160 daily transfers.

### Individual isolates from day-160 CAB cultures show diversity

That day-160 CAB populations attained substantially higher growth yields than day-5 populations (Figure 1D) is evidence of a phenotypic fitness change and likely accumulation of mutations. To assess the issue of mutation accumulation we sequenced the genomes of two isolates from lineage D and two isolates from lineage E day-160 CAB cultures. With the parent strain PAO1 as a reference we found between 151 and 167 single nucleotide polymorphisms in a given isolate. Both of the isolates from lineage E had a 6-kb deletion encompassing *pvdD* and *pvdJ*, genes required for synthesis of the iron siderophore pyoverdine (Table 1). Neither isolate from lineage D had this deletion, however both D isolates possess an internal gene duplication event that would render the pyoverdine peptidase synthase, *pvdI*, inactive (Table 1). The lineage E isolates had an amplification of a genomic region that included *nuh* of about 150 kb (two copies in one isolate and three in the other). One of the two lineage D isolates had two copies of a 90-kb *nuh*-containing region and the other isolate did not have amplification of a *nuh*-containing region (Table 1). As discussed above, mutations in *psdR* have been reported to sweep through populations grown on casein alone after just a few daily transfers and in fact we found *psdR* mutations in all four of the day-160 isolates. As expected, the *psdR* mutations were the same as those fixed in the day-50 populations (Table 1). Including the *psdR* mutations, there were nonsynonymous or in/del mutations in only 12 genes in all four of the sequenced isolates (Table 1, bold entities). Taken together, the genomic diversity and the improved population fitness suggest both that there are certain mutations that are selected in these conditions and also that there might be a division of labor among cells in these populations.

**Table 1.** Mutations in day-160 isolates present in at least one isolate from lineages D and E

| Locus<sup>a</sup> | Gene(s)<sup>b</sup> | Mutation | 160-day Isolate | SNP<sup>c</sup> | DNA coordinate(s) | Effect<sup>d</sup> | Gene description<sup>b</sup> |
| --- | --- | --- | --- | --- | --- | --- | --- |
| Multiple | PA0109-PA0192 | Gene duplication | D1 |  | 132441-222530 | 90.1 kb duplication (2 copies) | Contains two genes (encoding a purine nucleosidase and an adenosine deaminase) that when duplicated confer fast growth on adenosine (Toussaint, et al. 2017) |
|  | PA0117-PA0260 |  | E1 |  | 137061-291830 | 154.7 kb duplication (2 copies) |  |
|  | PA0122-PA0260 |  | E2 |  | 137061-290290 | 153.2 kb duplication (3 copies) |  |
| PA0143 | **nuh** | Intergenic SNP | D1, D2, E1, E2 | C>T | 163308 | Promoter mutation | Purine nucleosidase Nuh |
| PA0928 | **gacS** | Non-synonymous SNP | D1, D2, E1, E2 | C1420T | 101433 | G474S | Histidine kinase sensor-response regulator |
| PA1949 | **rbsR** | Non-synonymous SNP | D1, D2, E1, E2 | C32T | 2134424 | A11V | Ribose operon repressor RbsR |
| PA2338 |  | Non-synonymous SNP | E2 | A635G | 2582731 | Q212R | Probable binding protein component of ABC maltose/mannitol transporter |
|  |  | Insertion | D1, D2 | G856GC | 2582952 | R288 Frameshift |  |
| PA2399-PA2400 | **pvdD, pvdJ** | Deletion | E1, E2 |  | 2664201-2670360 | 6.2 kb deletion spanning 2 genes | Pyoverdine synthetase D, J |
| PA2402 | **pvdI** | (Internal) gene duplication | D1 |  | 2677291-2684990 | Internal 7.7 kb duplication (2 copies) | Pyoverdine peptide synthetase I |
|  |  |  | D2 |  | 2677291-2684220 | Internal 6.9 kb duplication (4 copies) |  |
| PA2492 | **mexT** | Non-synonymous SNP | D1, D2, E1, E2 | T887C | 2808355 | L296P | Transcriptional regulator MexT |
| PA2494 | **mexF** | Non-synonymous SNP | D1, D2, E1, E2 | G2069A | 2812077 | R690H | Multidrug efflux transporter MexF |
| PA2602<sup>e</sup> |  | Insertion | D2 | C338CG | 2945601 | R116 Frameshift | 3-mercaptopyruvate dioxygenase |
|  |  |  | E1, E2 | C338CGG | 2945601 |  |  |
| PA3205 |  | Insertion | E1 | TG81TCC | 3595466 | P31 Frameshift | Hypothetical protein |
|  |  | Insertion | D2 | TG81TGC | 3595466 |  |  |
| PA3488<sup>f</sup> | **tli5** | Insertion | E1 | A72AG | 3906126 | H28 Frameshift | T6SS-associated effector immunity protein |
|  |  | Deletion | D1, D2 | AG73A | 3906126 | G27 Frameshift |  |
|  |  | Insertion | E2 | C280CA | 3906334 | N96 Frameshift |  |
| PA3527<sup>g</sup> | **pyrC** | Intergenic insertion | D2, E1 | G>GC | 3947041 | Upstream mutation at 3947041 | Dihydroorotase |
| PA3535 | **eprS** | Stop gained | D1, D2, E1, E2 | C2491T | 3957397 | Q831Stop | Probable serine protease |
| PA3548 | **algI** | Intergenic SNP | D1, D2, E1, E2 | G>A | 3974147 | Upstream mutation at 3974147 | Alginate biosynthesis protein |
| PA3620 | **mutS** | Deletion | D1, D2, E1, E2 | TGC2067T | 4056590 | L690 Frameshift | DNA mismatch repair protein MutS |
| PA3626 |  | Non-synonymous SNP | E1, E2 | T232C | 4062198 | S78G | tRNA pseudouridine synthase D |
|  |  |  | D1 | T658C | 4061772 | S220G |  |
| PA3629 | **adhC** | Non-synonymous SNP | E1 | T800C | 4064188 | N267S | Alcohol dehydrogenase |
|  |  |  | D2 | T1033C | 4063955 | T345A |  |
| PA4344 |  | Non-synonymous SNP | E1, E2 | G449C | 4874042 | A150V | Probable hydrolase |
|  |  |  | D1 | T953C | 4873538 | E318G |  |
| PA4499 | **psdR** | Insertion | D1, D2 | A431AC | 5036675 | L146 Frameshift | Dipeptide regulator |
|  |  | Non-synonymous SNP | E1, E2 | T515C | 5036759 | V172A |  |
| PA4547 | **pilR** | Non-synonymous SNP | D1, D2, E1, E2 | T971C | 5095955 | L324P | Two-component response regulator PilR |
| PA4554 | **pilY1** | Deletion | E1, E2 | GC341G | 5101022 | N116 Frameshift | Type IV pilus assembly protein PilY1 |
|  |  | Stop gained | D1 | G962A | 5101644 | W321 Stop |  |
| PA4701<sup>h</sup> |  | Non-synonymous SNP | E1, E2 | T26C | 5280530 | L9P | Conserved hypothetical protein |
|  |  | Insertion | D1, D2 | A44AC | 5280548 | H17 Frameshift |  |
| PA5090<sup>i</sup> | **vgrG5** | Insertion | E1 | G1277GC | 5730568 | L427 Frameshift | T6SS-associated protein VgrG5 |
|  |  | Deletion | D2 | GC1277G | 5730568 | G426 Frameshift |  |

For inclusion in this list, the indicated gene(s) had mutations identified in one or both sequenced isolates from the two lineages.
Unless otherwise indicated, SNPs were identified by the presence of 90% or more variant reads at the indicated locus as compared to the PAO1 reference genome.
<sup>a</sup>Bold text indicates loci with SNPs or in/dels shared among all four isolates from the day-160 lineages with minimum read frequencies of 90% (except as in footnotes f - i).
<sup>b</sup>As described at pseudomonas.com (Windsor, et al. 2016).
<sup>c</sup>Nucleotide change and location within the coding sequence of the gene, unless the SNP is intergenic.
<sup>d</sup>Protein amino acid location and change resulting from the SNP or, in the case of intergenic SNPs, the genomic coordinate of the nucleotide change relative to the translation start codon.
<sup>e</sup>PA2602 SNP frequency was below the 90% variant reads cutoff for all four isolates, but present at notable frequency (89% variant reads) in lineage D isolate 2 and lineage E isolate 2, and in lineage E isolate 1 (80% variant reads).
<sup>f</sup>PA3488 SNP resulting in G27 frameshift was present in both isolates from lineage D supported by 94-95% variant reads, while the SNP resulting in H28 frameshift was present in lineage E isolate 1 with 79% variant reads; the N96 frameshift was found in both isolates from lineage E supported by 77 and 88% variant reads.
<sup>g</sup>PA3527 SNP frequency of 94% variant reads in lineage E isolate 1, and 84% in lineage D isolate 2.

### Validation of a population transcriptomics approach

Valuable information about the *P. aeruginosa* QS regulon has been gained by transcriptome analysis using QS gene deletion mutants (Schuster et al. 2003, Wagner et al. 2003, Chugani et al. 2012), but transcriptomics is a far from perfect tool and this is particularly true for identification of the complete set of QS-activated genes. Many genes in the *P. aeruginosa* QS regulon are co-regulated by other factors (Schuster et al. 2004). Some genes show delays in QS activation compared to other genes, and expression of other genes can be suppressed later in growth (Schuster et al. 2003). Previous investigations have shown that assessing the transcriptome during the transition between logarithmic phase and stationary phase (optical density of 2 in buffered LB) provides a reasonable census of QS-activated genes (Chugani et al. 2012). Thus, our analyses were with cells harvested from buffered LB at an optical density of 2.

Because we sought to define genes activated by QS, our comparison was to cells in which QS was inhibited. Such cells cannot grow in CAB by themselves (because they require QS for growth in this medium). Since our limited genomic sequencing analysis showed diversity among individuals within a lineage, we chose to perform transcriptomics on populations. This precluded the standard approach of generating QS mutants and comparing them to the wildtype. Rather we compared populations grown in buffered LB plus an AHL lactonase to populations grown with added AHLs (to control for the timing and magnitude of signal production by the populations). Previous publications have shown that addition of the *Bacillus* AHL lactonase AiiA to cultures can be employed to identify QS-activated genes in *P. aeruginosa* and other bacterial species (Feltner et al. 2016, Mellbye et al. 2016, Liao et al. 2018, Cruz et al. 2020). Unlike the growth results in CAB (Figure 1D), buffered LB growth of populations evolved for 160 days was indistinguishable from growth of populations evolved for only 5 days (Supplemental Figure S1). Addition of AiiA lactonase did not affect the growth of any of the *P. aeruginosa* populations and reduced AHLs to below detectable levels (<1 nM and 5 nM for 3OC12-HSL and C4-HSL, respectively).

We first compared the transcriptomes of lineage D and E populations taken from the fifth day of passage on CAB to each other. Populations were grown in buffered LB to an optical density of 2 with either added AHLs or added AiiA lactonase. Transcripts showing >2.8-fold higher levels in populations grown with added AHLs compared to populations grown with AiiA lactonase were considered QS activated. We identified 193 QS-activated genes in lineage D and 148 in lineage E (Table 2). There were 137 genes common to both lineages. These differences may reflect a combination of independent evolutionary trajectories and the vagaries of our transcriptome analysis. There is reason to believe that it may be the latter: most of the QS-activated genes identified in one, but not the other, lineage were minimally above the 2.8-fold threshold for classification as QS activated (Table 2). Several of these genes were in operons where other genes exceeded the cutoff. For example, 11 genes of the Hcp secretion island III (HSI-3) cluster of type VI secretion system genes (T6SS, PA2361-PA2371) in lineage D were classified as QS-activated, whereas only four of these genes in lineage E were classified as activated. Of the 137 QS-activated genes common to both lineages (Table 2), 126 (>90%) have previously been identified as such (Schuster et al. 2003, Chugani et al. 2012). For the 42 genes defined as core QS-activated genes (Chugani et al. 2012), 41 were QS-activated in both lineages D and E in the day-5 populations (Table 2). These results indicated that this transcriptomics approach would suffice to address our primary question: does long-term growth in CAB lead to a size reduction in the QS regulon?

**Table 2.**
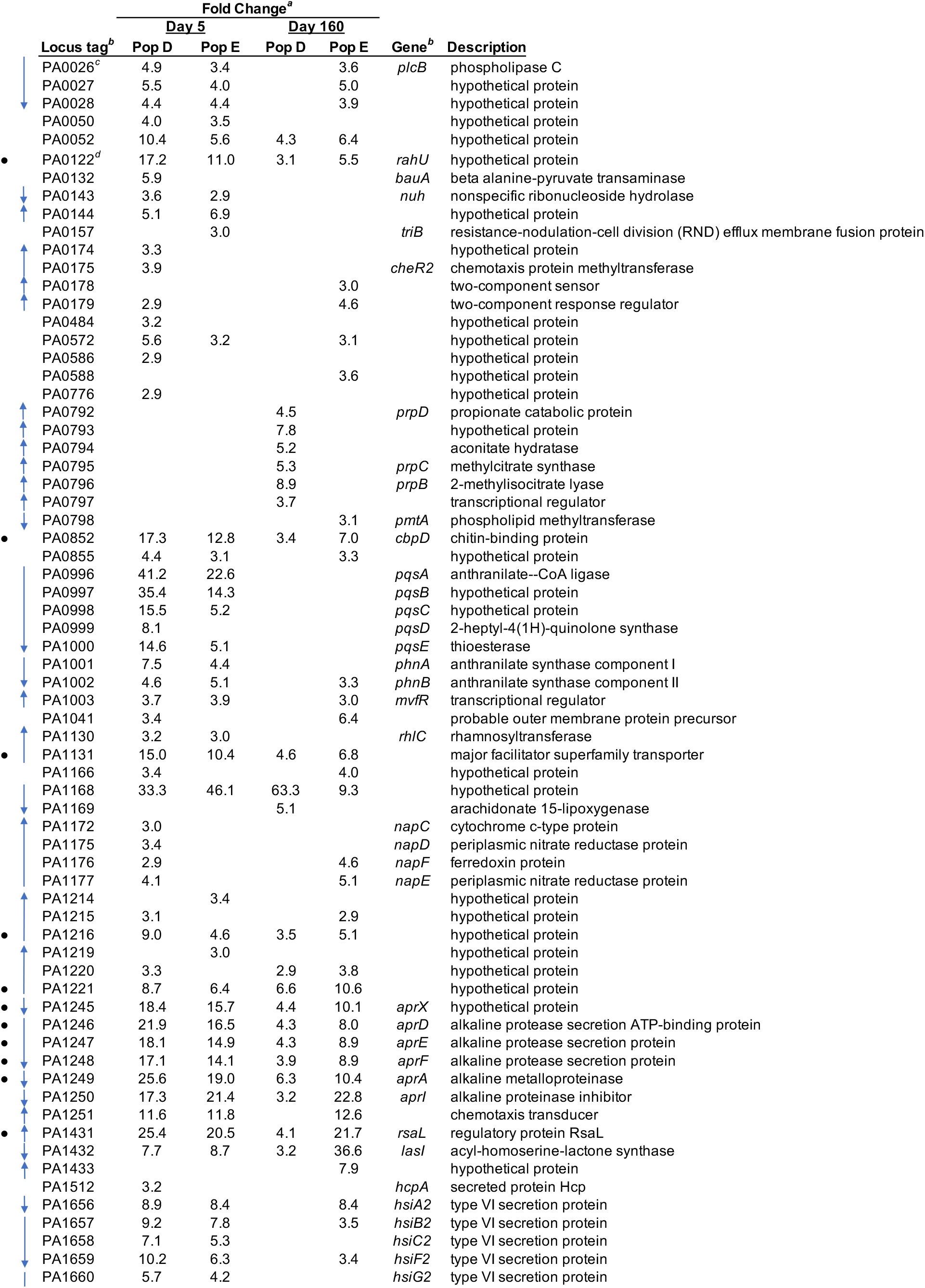
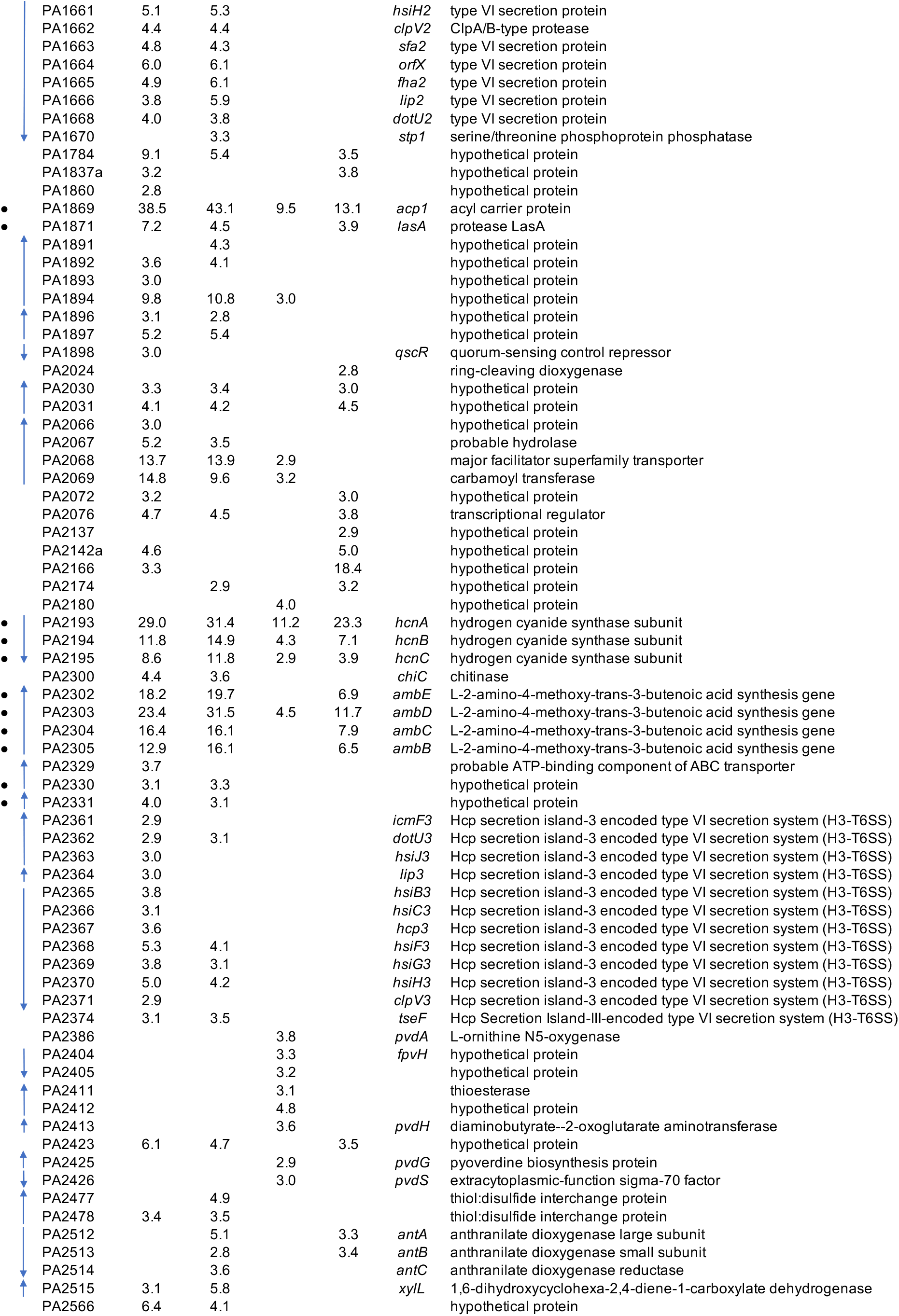
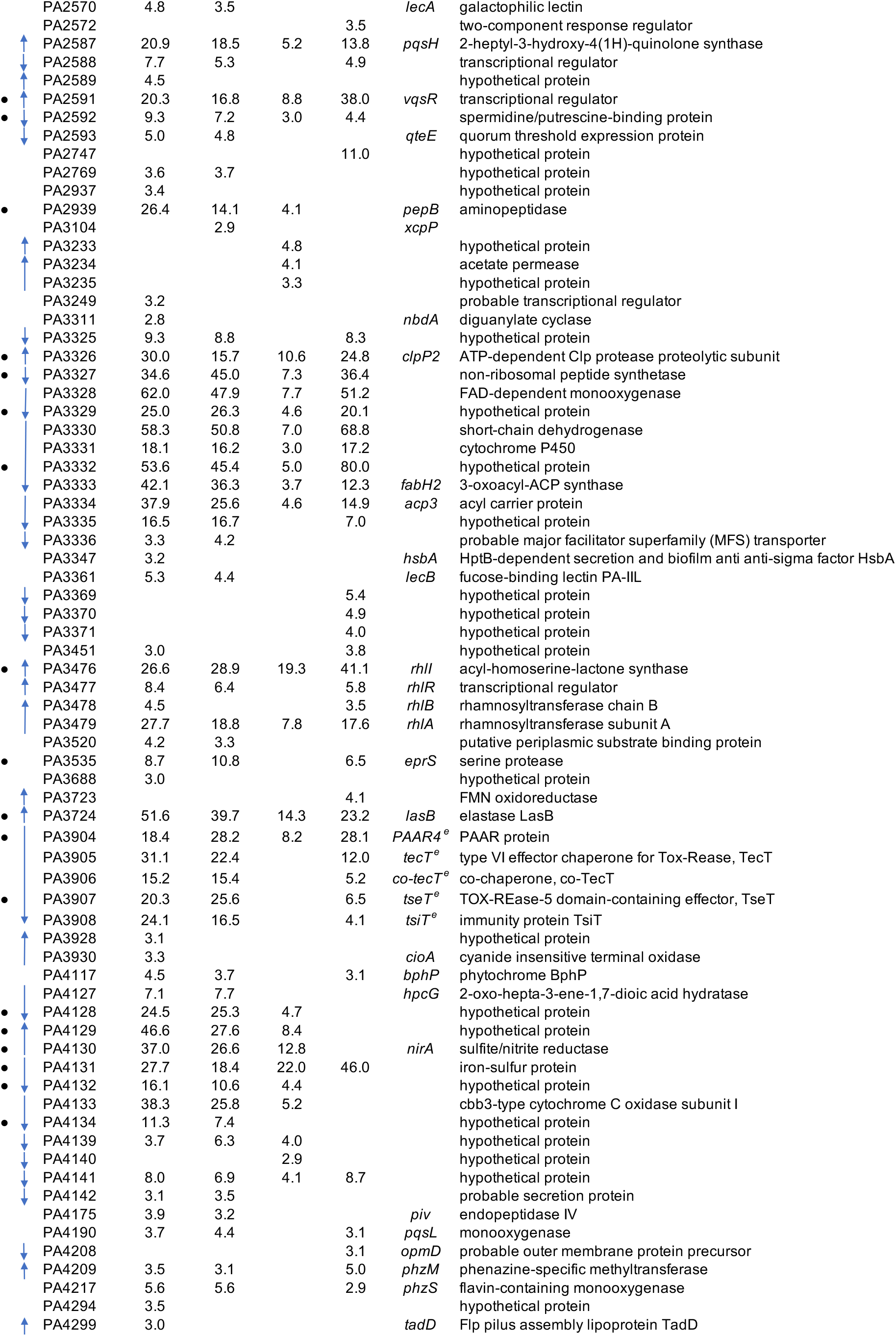
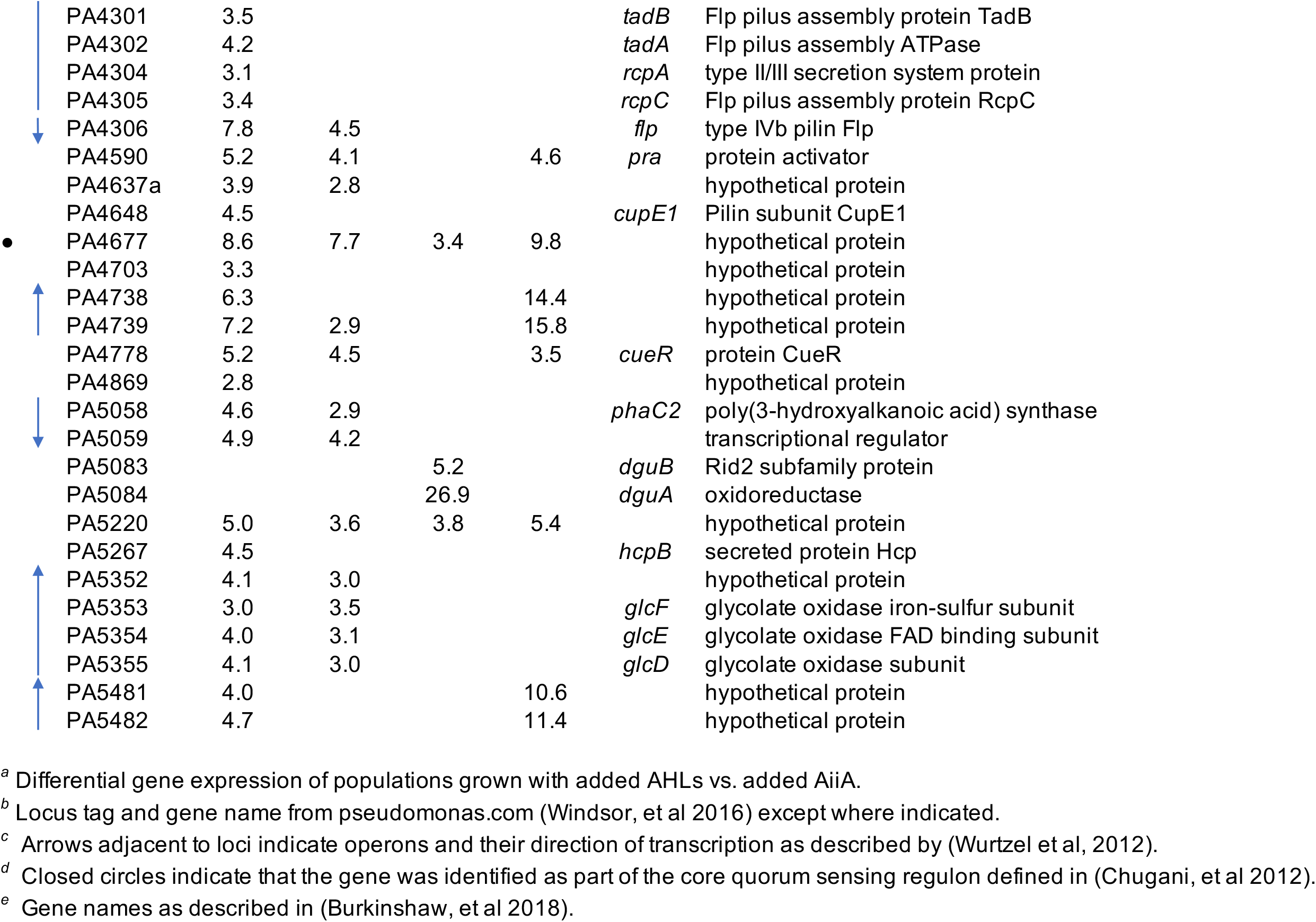
Quorum sensing-activated genes in day-5 and day-160 CAB populations

### Populations show a reduction in the QS regulon after 160 days in CAB

To determine the influence of long-term growth in CAB on the QS regulon, we performed RNA-seq on day-160 populations of lineages D and E grown in buffered LB with exogenously added AHLs and compared the transcriptomes to those grown with AiiA lactonase to identify QS-activated genes. We compared the QS-activated genes in day-160 populations to those in the respective day-5 populations (Table 2). The number of QS-activated genes in lineage D was reduced from 193 at day 5 to only 73 at day 160 (Figure 2A). For lineage E the number went from 148 to 108. There were a few genes that showed QS-activation in one, the other, or both day-160 populations but not in the day-5 populations. Without further experimentation we cannot speculate as to whether this is a result of the experimental approach or of biological significance. Regardless, there is a clear and obvious reduction in the number of genes in the QS regulon of day-160 populations.

**Figure 2.**
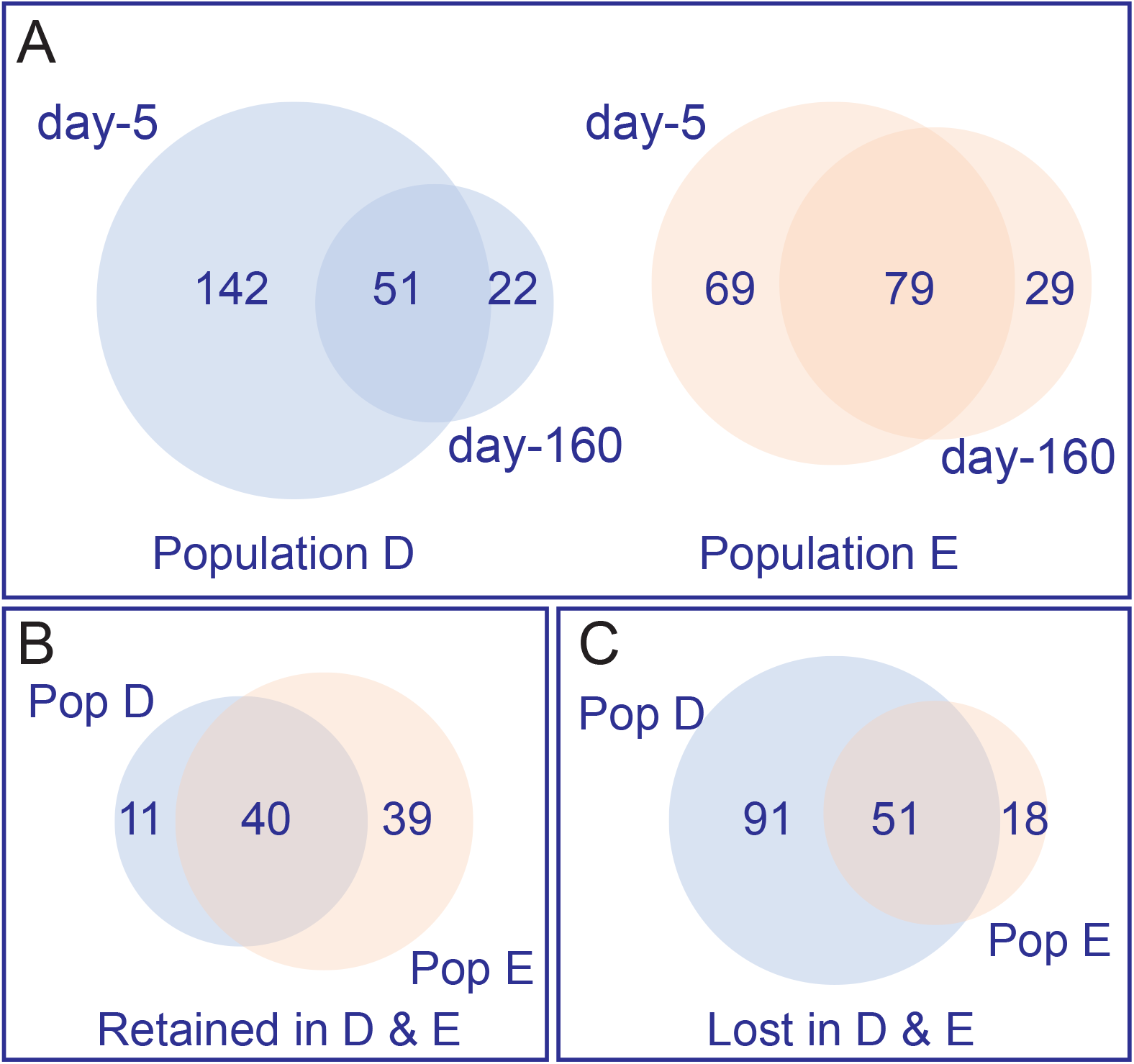
Many QS-activated genes in day-5 populations do not show QS activation in day-160 populations. (A) Venn diagrams showing the relationship between QS-controlled genes at day-5 vs. day-160 populations for lineages D (blue) and E (peach). (B) Venn diagram showing the overlap of genes that remain under QS-control (Retained) in both populations after 160-days of serial passage in CAB. (C) Venn diagram showing the overlap of genes that are no longer under QS-control (Lost) in both populations. Numbers in the Venn diagrams were determined using Venny (Oliveros 2015) and area calculated using the area-proportional Venn diagram plotter and editor found at http://apps.bioinforx.com/bxaf7c/app/venn/index.php. The numbers of genes in each category are indicated. Lists of genes shared in each category are in Table 2.

The loss of QS-gene activation after long-term growth in CAB could indicate a gene is expressed at levels similar to or greater than the QS-activated levels early in growth on CAB, or it could indicate a suppression of gene expression to levels similar to those in early populations grown in the presence of AiiA lactonase. To address this issue, we examined expression (Source Data File 1) of the 51 genes that showed QS-activation in both early populations, but not in both late populations. Only two of the 51 genes showed high levels of expression in day-160 populations, *nuh* and the adjacent gene PA0144 (Figure 3A). The high expression of *nuh* and PA0144 was expected from previous studies (Toussaint et al. 2017). The other 49 genes were expressed at very low levels from both 160-day populations in the presence of added AHLs (Figure 3B and Source Data File 1). This finding is consistent with the idea that the decreased size of the QS regulon after 160 days of growth on CAB is economical.

**Figure 3.**
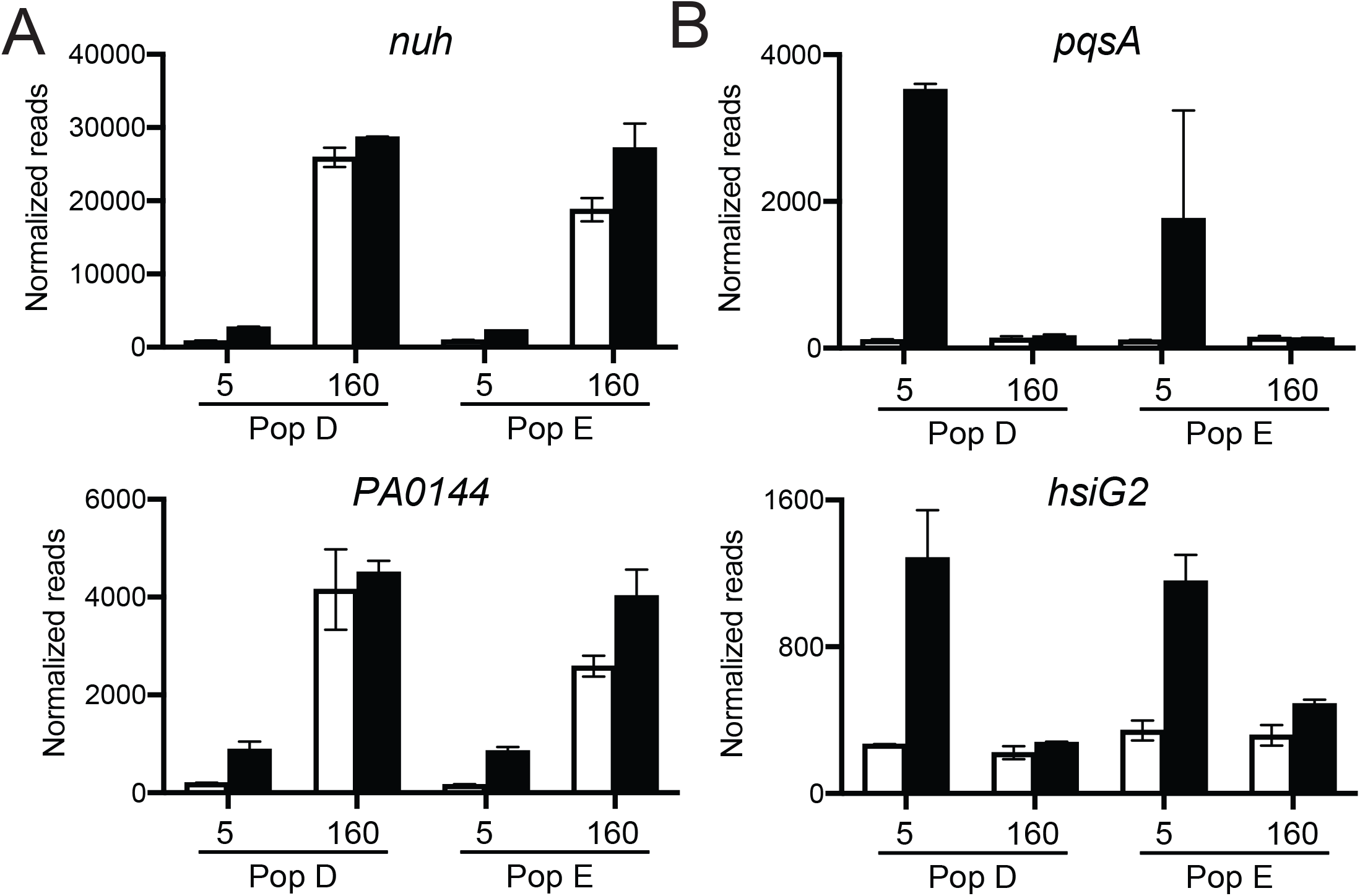
Normalized reads of select genes that are lost from QS control during serial passage in CAB for 160 days. (A) The only two lost genes with increased expression, *nuh* and PA0144, after 160 days of CAB passage. (B) Two representative lost genes, *pqsA* and *hsiG2*, with low expression levels in 160-day populations even in the presence of added AHLs. For all graphs white bars are AiiA-treated and black bars are plus AHL signals. Data are the mean normalized transcript counts of the two biological replicates for populations (D or E) passaged for 5 or 160 days in CAB; error bars represent the range. Normalized reads for all genes are provided in Source Data File 1.

### Characteristics of genes retained in the QS regulon after 160 days of growth in CAB

As expected, the genes for AHL synthesis, *lasI* and *rhlI*, are retained in the day-160 QS regulons of both lineages (40 genes in common are retained, Figure 2B). Genes for production of extracellular proteases are also retained, including the elastase gene, *lasB*, and the alkaline protease genes, *apr* genes (Table 2). Of note, not all genes coding for an extracellular protease are maintained in the regulon. For example, *piv*, which codes for an endopeptidase is no longer QS-controlled in both day-160 populations. Not all the genes retained in the QS regulon are obviously related to adenosine or casein digestion. For example, *hcnABC*, encoding a hydrogen cyanide synthase, remains QS-regulated; hydrogen cyanide has previously been shown to be important in policing of QS mutants (Wang et al. 2015, Yan et al. 2019). For other gene products, the relation to growth in CAB is unclear, and the significance of this finding is yet to be determined.

### Characteristics of genes lost from the QS regulon after 160 days of growth in CAB

There was a notable and obvious loss of genes related to Type VI secretion or Type VI-secreted products from the QS-regulons of both lineages (51 genes in common are lost, Figure 2C). For example, 9 of the 12 genes encoded by the Hcp secretion island II (HSI-2) cluster of T6SS genes (PA1656-PA1668) were eliminated from the QS regulons of lineages D and E by day 160, as were other T6SS-related genes throughout the genome. Continuous growth in the absence of interspecific bacterial competitors in CAB might minimize the usefulness of the Type VI secretion apparatus and secreted products that provide fitness by intoxicating local competitors (Schwarz et al. 2010, Hernandez et al. 2020). Three genes considered as core QS-activated genes (Chugani et al. 2012) were not activated in the day-160 populations of either lineage. All three (PA2330, PA2331, and PA4134) encode hypothetical proteins and showed substantial activation in day-5 populations but did not reach our threshold for activation in day-160 populations. Other such genes include a chitinase (*chiC*) and two lectins (*lecA/lecB*), that may be involved in nutrient acquisition in myriad environments, but not in CAB.

### Diverse routes to elimination of a gene from the QS regulon

The elimination of a given gene from the QS regulon by day 160 in CAB could have many possible explanations. For example, there can be a mutation in the promoter for that gene, there can be a delay in the QS activation of that gene such that it falls below the cutoff for activation at an OD of 2, there could be a mutation in a co-activator of that gene, or there could even be a mutation in that gene itself. Some of these possibilities have long-term consequences, and others can perhaps be more readily reversed if conditions change. To begin to address the question of how a gene identified as QS activated in day-5 populations might not be QS activated in day-160 populations, we examined promoter activity of the representative gene *pqsA* in our four day-160 clones used earlier for genome sequencing (two each, lineages D and E). We transformed the four isolates with a *pqsA-gfp* reporter plasmid and followed GFP fluorescence during growth in CAB. The two day-160 population E clones showed low, basal expression of the *pqsA* reporter over 16 hours of growth, indicating a loss of cell density-dependent expression associated with QS (Figure 4A). In contrast, only one of the two population D clones showed a similar loss in density-dependent *pqsA* expression, while the other showed *pqsA* expression consistent with QS activation, albeit delayed enough to prevent detection in our metatranscriptomic analysis.

**Figure 4.**
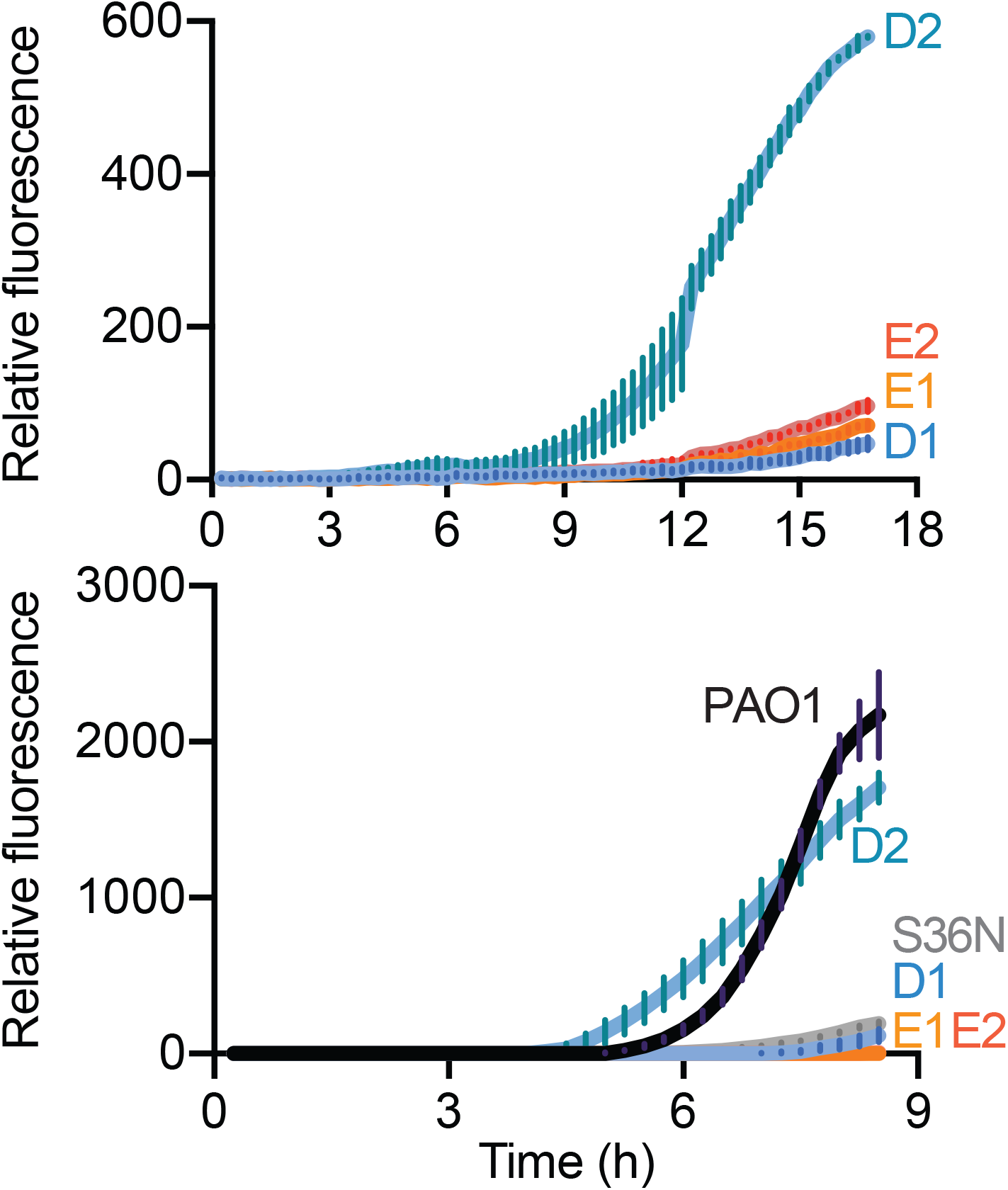
There are diverse routes to eliminate *pqsA* from the QS regulon. GFP fluorescence in isolates containing a P_*pqsA*_*-gfp* reporter plasmid, grown in 96-well plates in CAB (top) and buffered LB (bottom). The strains include isolates evolved for 160 days in CAB for population D (D1, D2) or E (E1, E2), the parent strain (PAO1), and a PqsR-variant strain (S36N). Plotted are the averaged relative fluorescence units over time of three technical replicates for two biological replicates; error bars represent standard deviations of means.

Our earlier genomic analyses identified an identical SNP in *pqsR* in both lineage E clones that codes for a S36N substitution. The *pqsR* gene (also called *mvfR*) codes for a PQS-responsive transcriptional activator of *pqsA* (Diggle et al. 2003, Deziel et al. 2005, Xiao et al. 2006). We reasoned that the PqsR S36N mutation could explain the loss of *pqsA* regulation by QS in our lineage E day-160 clones. To determine whether the PqsR S36N mutant is inactive and unable to induce expression of *pqsA*, we constructed a PAO1 *pqsR* S36N mutant and transformed both the mutant and PAO1 with the *pqsA-gfp* reporter. We monitored GFP-fluorescence during culture growth in buffered LB because *P. aeruginosa* PAO1 grows slowly in CAB during initial transfer from buffered LB. The S36N PAO1 mutant showed minimal GFP fluorescence compared to PAO1, similar to the lineage E clones (Figure 4B). For the sake of completeness, we also monitored GFP expression from the *pqsA-gfp* reporter in day-160 lineage D and E clones; the results were consistent with those from growth in CAB (Figure 4A). We conclude that the PqsR S36N mutation explains the loss of *pqsA* from QS regulation in lineage E. The varied, but delayed, expression of *pqsA* in the isolates from population D do not have as obvious of an explanation. Clearly, there are different evolutionary trajectories leading to loss of or delayed expression of genes like *pqsA in P. aeruginosa* populations during prolonged CAB growth.

## Discussion

*Escherichia coli* has been used as a model to investigate long-term evolution over a span of tens of thousands of generations in a specific growth environment, and this rich research area can be used as a framework for our studies of *P. aeruginosa*. For *E. coli* subjected to daily transfers to a fresh medium containing a single carbon and energy source, population fitness gains occur over time and certain mutations can sweep through a population (Lenski 2021). The sweeps result in a substantial gain in fitness. Fitness improvements are most dramatic early and become progressively smaller over time and generations. Multiple variants co-exist in a given population and evolutionary trajectories are complicated and varied (Good et al. 2017). It is notable that, unlike *E. coli*, our experimental subject, *P. aeruginosa*, is notorious for undergoing genetic changes during routine laboratory maintenance. In fact, strain PAO1 cultures that have been maintained in different laboratories show significant different genotypic and phenotypic characteristics (LoVullo and Schweizer 2020).

Compared to *E. coli*, much less is known about evolution of *P. aeruginosa* during adaptation to a constant environment. There is some information about adaptation to environments where *P. aeruginosa* QS is required for growth (Schuster et al. 2013), and examples of such genetic adaptations are present in all isolates passaged for 160 days in CAB (Table 1). In a relatively short time during QS-dependent growth on casein, a mutation in *psdR*, which codes for a repressor of a small-peptide uptake system, sweeps through the population and results in a substantial fitness gain reflected by more rapid growth in this environment (Asfahl et al. 2015). Inactivating mutations in the global regulator gene *mexT* or *mexEF-oprN* efflux pump genes also improve growth on casein, at least in part, by increasing activity of the Rhl QS system (Oshri et al. 2018, Kostylev et al. 2019). During QS-dependent growth on adenosine, a poor energy source for *P. aeruginosa* PAO1, amplification of a genomic region containing the adenosine hydrolase gene *nuh* arises and results in a substantial fitness advantage (Toussaint et al. 2017). Growth on casein requires the QS-induced public extracellular protease elastase (LasB), and after *psdR* sweeps through the population, LasR QS mutants emerge. These mutants, which do not activate any of the many genes in the QS regulon, have a negative frequency-dependent fitness advantage over their parents during growth on casein (Diggle et al. 2007, Sandoz et al. 2007, Schuster et al. 2013). QS-dependent growth on adenosine requires the periplasmic *nuh* product, purine hydrolase, a private good. In short-term evolution experiments (30 days, about 200 generations) on casein and adenosine together, QS mutants are constrained. They can benefit from proteases produced by their parents but they have lost access to adenosine (Dandekar et al. 2012). Together these adaptations to QS-dependent growth suggest plasticity in QS regulons, and particularly during growth under strong selective pressures.

In any given environment QS may be important for fitness as a result of activation of only a small number of genes. A fitness cost would be incurred by expression of the many other QS-activated genes. One solution to this fitness problem might be a reduction in QS activation of a large fraction of the QS regulon. There are several issues related to such a possibility. How might this be facilitated mechanistically? Might such a reduction in any given isolate reflect its ecological history? Would this provide a possible benefit in that perhaps a silenced gene could be returned to the QS regulon if ecological conditions are altered? The gene pool can remain intact when fitness is a result of rewiring gene regulation. There is circumstantial evidence to support the idea that the QS regulon might reflect the ecological history of a given isolate. In a limited comparison of QS-activated genes in seven isolates from different habitats, Chugani and colleagues found diversity in the regulons of QS-activated genes among various isolates (Chugani et al. 2012).

We approached the question of whether reductions in the QS regulon might reflect environmental conditions by executing a long-term evolution experiment with the well-studied *P. aeruginosa* strain PAO1 grown in a specific medium where only a fraction of the genes activated by QS would provide an obvious benefit to the population. To facilitate our experiments, we transferred populations daily in a medium containing both casein and adenosine (CAB) as the sources of carbon and energy. As expected, LasR QS mutants were constrained during *P. aeruginosa* PAO1 growth in CAB. By screening for protease-negative mutants we showed LasR mutants were constrained in all five of the lineages we maintained for at least 50 days. By day 80, LasR mutants had broken through in two of the five lineages (Figure 1B).

We subjected two of the three populations that continued to constrain LasR mutants at day 160 to further study. Similar to *E. coli*, fitness improvements were most dramatic early (by day 50), and became progressively smaller over time (Figure 1D). In fact, *psdR* mutations arose in both populations sometime between day 5 and day 50. It was also during this period that populations acquired the ability to grow rapidly on adenosine (Figure 1C): there was both genotypic and phenotypic heterogeneity within each population. This heterogeneity suggests an interesting possibility, and it creates an experimental dilemma. The possibility is that there might not only be competition for resources among the individuals in the population, but there might also be cooperation through a division of labor to enhance resource utilization. We have not yet addressed this possibility. The dilemma is that one cannot expect to learn about the breadth of the QS regulon by isolating a small number of individuals, creating QS mutants, and comparing transcriptomes of mutants to parents. To address this problem, we devised a method to interfere with population-level QS by using an AHL lactonase. We call this method QS meta-transcriptome gene analysis. Purified AHL lactonase has been employed to study QS gene activation in bacterial pure cultures (Feltner et al. 2016, Mellbye et al. 2016, Liao et al. 2018, Cruz et al. 2020). QS meta-transcriptome gene analysis extends this method and allows for studies of gene regulation in heterogeneous populations, even mixed-species populations.

Our QS meta-transcriptomic analysis revealed marked reductions in the number of genes activated by QS in day-160 populations compared to day-5 populations. In one population (D) the number of QS activated genes at day 160 was only 26% of the number at day 5, and in the other population (E) it was 53%. One way to eliminate a gene from the QS regulon is by deletion, but our genome sequencing of two isolates from each population indicated this was not a common event. Rather, it appeared that many QS regulon genes were intact and simply remained under our arbitrary threshold (2.8-fold) at our arbitrary point in growth. This could result from a number of mechanisms including delay in QS gene activation, modification of a co-regulatory pathway, and subtle changes in levels of other transcriptional regulators in cells. Regardless of the mechanism, this seems to represent a flexible solution to reducing the cost of QS when *P. aeruginosa* populations thrive in an environment where only a few QS activated genes provide a benefit. Although as yet untested, we hypothesize that modifying the environment in one way or another to involve QS activities other than protease or adenosine metabolism can further alter the regulon bringing genes back under QS control.

To begin to understand ways in which *P. aeruginosa* reduced its QS regulon during long-term growth in CAB we focused on *pqsA*, a gene that showed greater than 10-fold QS activation in either day-5 population and no activation in either day-160 population (Table 2, Figure 3B). *pqsA*, the first gene in the *pqs* operon, is required for production of the *Pseudomonas* Quinolone Signal (PQS). The *pqsA* promoter is activated by the PqsR transcription factor together with PQS, and *pqsR* transcription is activated by LasR. We found a point mutation in the two day-160 lineage E isolates, and this mutation was in fact fixed in the day-160 population. This presents a solution to the question of how the *pqs* operon is removed from QS activation and silent. In this solution, the pseudogene leaves an inactive copy of *pqsR* in the genome and pseudogenes can be repaired. The two population D isolates showed different *pqsA* expression patterns in pure culture. One exhibited a low level of *pqsA* expression and the other showed delayed expression in buffered LB. Neither had a *pqsR* mutation. This population has developed a different strategy, as yet unknown, to release *pqsA* expression from QS activation. This finding is consistent with our other results indicating the populations are heterogeneous.

That genes can be lost from QS-control under the specific conditions of our experiment supports the view that assessing the QS regulon of a *P. aeruginosa* isolate might provide information about the ecological history of that isolate. We note that some of the differences between day-160 lineages and between day 5 and day 160 in a given lineage might be due to the noise inherent in RNA-seq analyses. Nevertheless, during the course of our experiment, there was a reduction in the number of genes activated by QS. The increased population fitness observed between day 50 and day 160 in our metapopulations (Figure 1D) at least in part result from this economization in the size of the QS regulon.

## MATERIALS AND METHODS

### Bacterial strains, plasmids and growth conditions

Strains, plasmids and primers used for the study are listed in Supplemental Index Tables S1 and S2. To create the PqsR S36N variant strain we used a homologous recombination-based, two-step allelic exchange approach as described previously (Kostylev et al. 2019). Bacteria were grown in either lysogeny broth [modified to use 0.5% NaCl (Miller 1972)] buffered with 50 mM 3-(N-morpholino)propanesulfonic acid at pH 7.0 (buffered LB) or the minimal medium described in (Kim and Harwood 1991) with 0.25% caseinate and 0.75% adenosine (CAB) (Dandekar et al. 2012) or 1% adenosine-only broth (Figure 1C). All broth cultures were grown at 37°C with shaking (250 rpm) in 18 mm tubes, or in 200 μl volumes in 96-well plates where indicated. When required, gentamicin was added to the medium (10 μg/ml *E. coli*, 100 μg/ml *P. aeruginosa*).

### Long-term growth experiments and phenotypes

Evolved populations were derived from wild-type *P. aeruginosa* strain PAO1 as diagrammed in Figure 1. Five individual colonies were used to inoculate buffered LB. Overnight cultures were used to inoculate tubes containing 4-ml CAB (1% inoculum vol/vol). After a 24-h incubation, 100 µl of culture was used to inoculate a fresh tube of CAB. This process continued for 160 days. One-ml volumes of each population were collected at days 5, 20, 50, 80, and 160 and stored as frozen glycerol (25% vol/vol) stocks at -80°C. To screen for protease-negative isolates (Figure 1B) we plated onto LB-agar and then patched 100 individual colonies onto skim milk agar plates (Sandoz et al. 2007). Colonies that lacked a zone of clearing on the milk plates were scored as protease-negative. To look for *psdR* mutations, scrapings of day-5 and day-50 population frozen glycerol stocks were used to inoculate buffered LB. After overnight incubation, cells were pelleted by centrifugation and genomic DNA was extracted using the Gentra Puregene Yeast/Bacteria kit (Qiagen, Germantown, MD). The purified DNA was used as template for PCR (*psdR* primers provided in Supplemental Index Table S2) and the PCR product Sanger sequenced. For growth in 1% adenosine-only broth (Figure 1C), scrapings of day-5 and day-50 population frozen glycerol stocks or individual colonies of PAO1 were used to inoculate buffered LB and incubated overnight. Overnight cultures were back diluted in 1% adenosine-only broth to an OD_600_ of 0.01 and growth monitored by absorbance at 600 nm. To determine cell yields on CAB, scrapings of population frozen glycerol stocks were used to inoculate buffered LB and incubated overnight. A 3-ml CAB culture was then inoculated with 30 μl of overnight culture, incubated at 37°C with shaking for 18 h, and then serially diluted for plating on LB agar to determine the number of colony forming units.

### Genome sequencing and variant analysis

Scrapings of day-160 population frozen glycerol stocks were streaked for isolation on LB agar plates. After an overnight incubation, two individual colonies from population D and two from population E were used to inoculate buffered LB. After overnight growth, genomic DNA was extracted from cells and purified by using the Gentra Puregene Yeast/Bacteria kit (Qiagen, Germantown, MD). The purified DNA was used to construct paired-end 2×150 bp libraries for sequencing on an Illumina MiSeq (San Diego, CA). Reads were aligned to the PAO1 reference genome (accession NC_002516) using the genomics software StrandNGS version 3.3.1 (Strand Life Sciences, Bangalore, India). Variant analysis was performed with StrandNGS SNP, CNV and SV pipelines. We defined a SNP as having a variant read frequency of >90% and an indel as being <100 bp. Previously, we sequenced the genomes of two individual isolates of our laboratory MPAO1 strain (SAMN06689578, SAMN09671539), which we used as comparisons for SNPs calls and copy number/indel analyses.

### Purification of AiiA lactonase

The AiiA AHL lactonase enzyme was purified as a maltose-binding protein (MalE) fusion from *E. coli* cells as described elsewhere (Thomas and Fast 2011) except that 0.2 mM CoCl2 was substituted for ZnCl2 and the MalE fusion was not removed by TEV cleavage. Purified AiiA was concentrated to 10 mg/ml in protein buffer [20 mM Tris-HCl, 100 mM NaCl, pH 7.4 with 10% glycerol (vol/vol)], and stored at -80°C until use. AiiA was added to cultures at a final concentration of 100 μg/ml, which was sufficient to reduce 3OC12- and C4-HSL levels to below detection in culture extracts as measured by bioassay (Schaefer et al. 2000).

### RNA-seq analyses

Scrapings from glycerol stocks of populations D and E day-5 and day-160 passages were used to inoculate into 3-ml of buffered LB. After an overnight incubation, cells from these cultures were used to inoculate 3-ml of buffered LB with a starting optical density (OD_600_) of 0.01. These cells were grown to logarithmic phase (OD_600_ 0.1-0.4) and then used to inoculate a 250-ml baffled flask containing 25-ml of buffered LB (starting OD_600_ 0.005) and one of the following: AiiA lactonase (100 μg/ml), AHLs (2 μM 3OC12-HSL and 10 μM C4-HSL final concentrations), or no additions. Two biological replicates were prepared for each sample condition. In early stationary phase (OD_600_ 2), AHL signals were solvent extracted from 5-ml of culture and concentrations were determined by bioassay as described previously (Schaefer et al. 2000) to confirm AiiA-lactonase activity. At the same time, cells from 2-ml of culture were pelleted by centrifugation, preserved in RNAprotect bacteria reagent (Qiagen, Germantown MD), and stored at -80°C, followed by RNA extraction as described previously (Cruz et al. 2020).

RNA-seq library preparation, bacterial Ribo-Zero rRNA depletion (Illumina, San Diego, CA), and sequencing was performed by Genewiz (South Plainfield, NJ). Library samples were divided into two separate HiSeq3000 runs, each with paired-end 2×150 bp read lengths. Trim Galore! (Babraham Bioinformatics, Cambridge UK) was used to trim adapters prior to alignment against the PAO1 reference genome (accession NC_002516) using StrandNGS version 3.3.1 (Strand Life Sciences, Bangalore, India). DESeq2 (Love et al. 2014) was used for differential expression analysis, using the Benjamini-Hochberg adjustment for multiple comparisons and a false-discovery rate α = 0.05. Samples from each treatment regimen were grouped by individual population for the DESeq2 differential expression analyses (i.e., the three treatments for lineage D day-5 populations described above were compared separately from the day-160 population treatments, and likewise for lineage E populations). QS-activated regulons were determined by comparing added AHLs vs. AiiA-treated samples for a given population (D or E) and passaging time (5-day or 160-day), and imposing a 2.8-fold minimal fold-change threshold. We elected to use the added AHLs treatment as the QS-induced treatment, rather than the no additions treatment where AHL signals accumulate during culture growth, to control for the timing and magnitude of AHL levels. However, the no additions treatment was included in the DESeq2 differential expression analyses. Normalized transcript counts (Source Data File 1 and Figure 3) were obtained by taking the inverse log of the regularized log transformation (rlog) function in DESeq2 (Love et al. 2014). Note that because there are two phenazine biosynthesis *phz* operons in the PAO1 genome with high levels of sequence identity, it is not possible to differentiate from which operon the transcripts derived, as the StrandNGS software assigns reads that match multiple locations (<5 matches) to the earliest match location in the genome. Thus, although phenazine synthesis is QS-regulated, we excluded *phzA1-G1* and *phzA2-G2* from subsequent analyses of our differential expression data.

### QS gene reporter activity during growth

We created a promoter probe fusion for *pqsA* and a promoterless *gfp* control (Supplemental Index Table S2) by using PCR and *E. coli*-mediated DNA assembly (Kostylev et al. 2015). Plasmids were used to electrotransform *P. aeruginosa* strains. Single transformant colonies harboring pBBR-P*pqsA*-*gfp* or pBBR-*gfp* were used to inoculate buffered LB supplemented with 100 µg/ml gentamicin. Overnight cultures were back diluted to OD_600_ 0.01, grown to mid-log phase (OD_600_ 0.1-0.4), diluted to an initial OD_600_ of 0.01 in CAB or buffered LB as indicated, and dispensed in 200 μl volumes into a 96-well microtiter dish plate. Fluorescence (excitation 485 nm, emission 535 nm) and absorbance (OD_600_) were measured every 15 min for 14 h using a Biotek Synergy H1 microplate reader. Background fluorescence was determined using the promoterless pBBR-*gfp* control.

### Sequence data deposition

Raw sequencing reads for variant analyses of the two individual isolates from each of the day-160 lineages (BioSamples SAMN16400702-SAMN16400711) were deposited in the NCBI Sequence Read Archive under BioProject PRJNA667949. Raw sequencing reads and count matrices for transcriptome analyses of CAB-evolved populations D (BioSamples SAMN16400698, SAMN16400699) and E (BioSamples SAMN16400700, SAMN16400701) were deposited under GEO accession GSE176411 as well.

## ACKNOWLEDGEMENTS

This work was supported by US Public Health Service (USPHS) Grants K08AI102921 and R01GM125714 (to A.A.D.) and R01GM059026 and R35GM136218 (to E.P.G.). A.A.D. was also supported by the Burroughs-Wellcome Fund (Grant 1012253). E.P.G. and A.L.S. were also supported by the Genomic Science Program, US Department of Energy, Office of Science, Biological and Environmental Research as part of the Plant-Microbe Interfaces Science Focus Area (pmi.ornl.gov). Oak Ridge National Laboratory is managed by University of Tennessee-Battelle LLC for the US Department of Energy under Contract DE-AC05-00227525. K.L.A. was supported by a Postdoctoral Fellowship from the Cystic Fibrosis Foundation (ASFAHL19F0). We acknowledge core support from USPHS Grant P30DK089507 and the Cystic Fibrosis Foundation (Grants SINGH15R0 and R565 CR11).

## Notes

### Competing Interest Statement

The authors have declared no competing interest.

