## Supplemental Tables 1, 2 and Supplemental Figure 1 for "Evolution of the quorum sensing regulon in cooperating populations of *Pseudomonas aeruginosa*"

Table S1. Strains used in this study

| Bacterial strain | Description | Source |
|---|---|---|
| ***P. aeruginosa*** | | |
| MPAO1 | Wild-type *P. aeruginosa* | (1) |
| D1 | Isolated from Population D after 160 passages in CAB | This work |
| D2 | Isolated from Population D after 160 passages in CAB | This work |
| E1 | Isolated from Population E after 160 passages in CAB | This work |
| E2 | Isolated from Population E after 160 passages in CAB | This work |
| NES24 | PqsR (MvfR) S36N variant in MPAO1 background | This work |
| ***E. coli*** | | |
| DH5α | *fhuA2 Δ(argF-lacZ)U169 phoA glnV44 Φ80 Δ(lacZ)M15 gyrA96 recA1 relA1 endA1 thi-1 hsdR17* | Invitrogen |
| S17-1 | *thi pr, hdsR hdsM+ rec,* RP4-2 (Tc::Mu Km::Tn7) | (2) |

Table S2. Plasmids and primers used in this study

| Plasmids | Relevant characteristics | Source |
|---|---|---|
| pMAL-t-*aiiA* | Maltose binding protein-AiiA fusion vector, Ap$^R$ | (3) |
| pPROBE-GT | Source for multiple cloning site (mcs) *gfp* fusion, Gm$^R$ | (4) |
| pBBR1MCS-5 | Broad-host range plasmid, Gm$^R$ | (5) |
| pBBR-*gfp* | Promoterless *gfp* transcriptional reporter; pBBR1MCS with mcs, *gfp* fusion from pPROBE-GT, Gm$^R$ | This work |
| pP$_{pqsA}$-*gfp* | pBBR-*gfp* with -429 to 0 of the *pqsA* promoter, Gm$^R$ | This work |
| pEXG2 | Allelic exchange vector with pBR origin, *sacB*, Gm$^R$ | (6) |
| pNS12 | pEXG2 containing sequences to create the PqsR S36N variant allelic exchange vector, Gm$^R$ | This work |

| Primer | DNA sequence (5' to 3') | Description |
|---|---|---|
| psdR_F | GGGTTCGGGTAGTTCATC | *psdR* SNP detection |
| psdR_R | GTTTGCCTGACAGGATGG | *psdR* SNP detection |
| mcsGFPpBBR_F | ATCGGTGCGGGCCTCTTCGCTATTACGCCAGCAAGCT TGCATGCCTGCAGGTC | pBBR-gfp construction |
| GFP_pBBR_R | CTAAAGGGAACAAAAGCTGGGTACCCTATTTGTATAGT TCATCCATGCCATGTGTAATCC | pBBR-gfp construction |
| vec_pBBRGFP_F | ATGGCATGGATGAACTATACAAATAGGGTACCCAGCTT TTGTTCCCTTTAGTGAG | pBBR-gfp construction |
| vec_pBBRGFP_R | ATCCTCTAGAGTCGACCTGCAGGCATGCAAGCTTGCT GGCGTAATAGCGAAGAGG | pBBR-gfp construction |
| PpqsA_pBBR_F | GGGCCTCTTCGCTATTACGCCAGCAAGATGCCGTCGC CCCCTTGGAG | pP$_{pqsA}$-*gfp* construction |
| PpqsA_pBBR_R | ATCCTCTAGAGTCGACCTGCAGGCATGCAAGCATGAC AGAACGTTCCCTCTTCAGC | pP$_{pqsA}$-*gfp* construction |
| vec_pBBRpqsA_F | TATCGCTGAAGAGGGAACGTTCTGTCATGCTTGCATGC CTGCAGGTCGAC | pP$_{pqsA}$-*gfp* construction |
| vec_pBBRpqsA_R | TGGGCTCCAAGGGGGCGACGGCATCTTGCTGGCGTA ATAGCGAAGAGG | pP$_{pqsA}$-*gfp* construction |
| pqsR_SNP1_F | ATTGCAACTGGTCTATTTTCCTCTTATGCTGGTTGCCG AAACGGGCCATC | pNS12 construction |
| pqsR_SNP2_R | TGCTGACCGCCGAGtTGACCGCGGTGTG | pNS12 construction |
| pqsR_SNP3_F | TCGCACACCGCGGTCAaCTCGGCGGTCAGCAACCTG | pNS12 construction |
| pqsR_SNP4_R | TCCTTTTATGATTTTCTATCAAACAATTCCATCCCGAGT CGATTCTCACCACCCACGGCCA | pNS12 construction |
| pEXG2-pqsRS36N_F | GCCGTGGGTGGTGAGAATCGACTCGGGATGGAATTGT TTGATAGAAAATCATAAAAGGATTTG | pNS12 construction |
| pEXG2-pqsRS36N_R | TCGCTGGAGATGGCCCGTTTCGGCAACCAGCATAAGA GGAAAATAGACCAG | pNS12 construction |

Ap$^R$, resistant to ampicillin; Gm$^R$, resistant to gentamicin

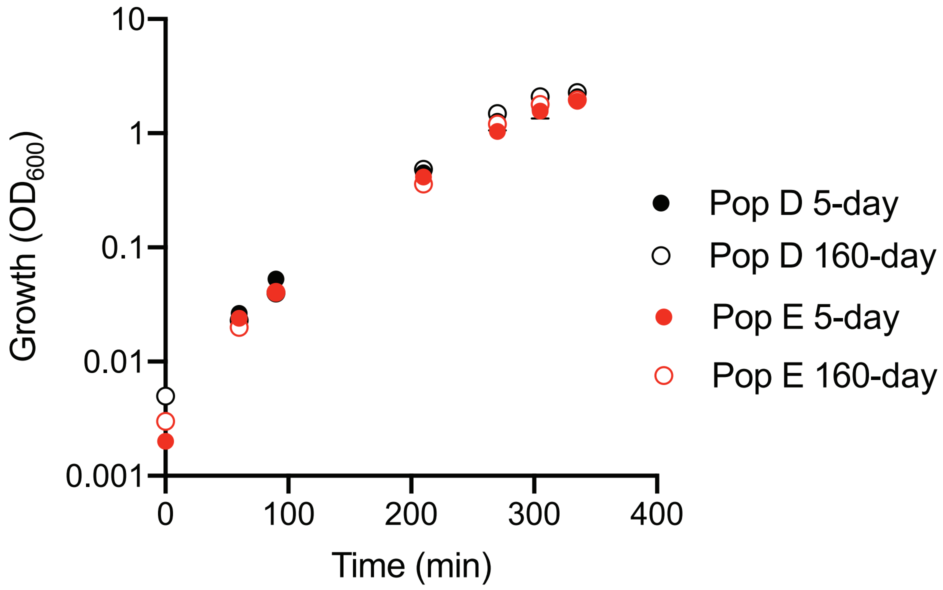


Figure S1. Growth of populations from 160-day CAB cultures (filled symbols) show similar growth in buffered LB as do populations from 5-day CAB cultures (open symbols). Preparation of inocula and growth conditions in 250-ml flasks were as described in the RNA-Seq Analyses section of the Materials and Methods. The data are means of two biological replicates; error bars indicate the ranges.
